# SARS-CoV-2 reactive T cells in uninfected individuals are likely expanded by beta-coronaviruses

**DOI:** 10.1101/2020.07.01.182741

**Authors:** Ulrik Stervbo, Sven Rahmann, Toralf Roch, Timm H. Westhof, Nina Babel

**Affiliations:** Center for Translational Medicine, University Hospital Marien Hospital Herne, Ruhr-University Bochum, Germany; Berlin-Brandenburg Center for Regenerative Therapies, and Institute of Medical Immunology, Charité – Universitätsmedizin Berlin, Corporate Member of Freie Universität Berlin, Humboldt-Universität zu Berlin, and Berlin Institute of Health, Berlin, Germany; Genome Informatics, Institute of Human Genetics, University of Duisburg-Essen, Germany

**Author notes:** Equal contribution (Ulrik Stervbo, Sven Rahmann). Contact: Ulrik Stervbo, Sven Rahmann.

## Abstract

The current pandemic is caused by the SARS-CoV-2 virus and large progress in understanding the pathology of the virus has been made since its emergence in late 2019. Several reports indicate short lasting immunity against endemic coronaviruses, which contrasts repeated reports that biobanked venous blood contains SARS-CoV-2 reactive T cells even before the outbreak in Wuhan. This suggests there exists a preformed T cell memory in individuals not exposed to the pandemic virus. Given the similarity of SARS-CoV-2 to other members of the Coronaviridae family, the endemic coronaviruses appear likely candidates to generate this T cell memory. However, given the apparent poor immunological memory created by the endemic coronaviruses, other immunity against other common pathogens might offer an alternative explanation. Here, we utilize a combination of epitope prediction and similarity to common human pathogens to identify potential sources of the SARS-CoV-2 T cell memory. We find that no common human virus, other than beta-coronaviruses, can explain the pre-existing SARS-CoV-2 reactive T cells in uninfected individuals. Our study suggests OC43 and HKU1 are the most likely pathogens giving rise to SARS-CoV-2 preformed immunity.

## Introduction

The current coronavirus disease 2019 (COVID-19) pandemic is caused by the severe acute respiratory syndrome coronavirus-2 (SARS-CoV-2) (1) with devastating consequences. SARS-CoV-2 infections have a broad spectrum of manifestations, ranging from asymptomatic to severe pneumonia and acute respiratory distress syndrome (2). The reason for this broad range is unclear, but markedly decreased immune cell numbers (3,4), together with cytokine storm (5), and dysregulation of lung infiltrating immune cells (6–9) have been associated with critical COVID-19 manifestations. The virulence of SARS-CoV-2 is in contrast to the four endemic coronaviruses (OC43, HKU1, 229E, NL63), which are responsible for the common cold with usually mild symptoms (10).

T cells recognise peptides presented in the context of the human leukocyte antigen (HLA) class I and class II molecules. Peptides presented on the class I HLAs are generally recognised by CD8^+^ cytotoxic T cells, while CD4^+^ T helper cells recognise peptides bound on the HLA class II molecule. T cells are known to be cross reactive (11,12), that is, a single T cell can recognise similar peptides derived from different pathogens presented by HLA molecules.

The SARS-CoV-2 virus elicits a T cells response during the infection (13). However, mounting evidence suggests that also uninfected individuals are capable of responding to peptides derived from the S-protein of the SARS-CoV-2 virus (13–17), indicating a pre-existing immunity to the SARS-CoV-2 S protein. A single influenza epitope has been identified by comparison of T cell receptors (18), but the original pathogenic source of this pre-formed T cell memory is generally unclear.

Given the high sequence similarity, the endemic coronaviruses are likely inducers of the preexisting immunity. However, immunity to these coronaviruses appears short-lived as antibody titers return to baseline levels at 4-12 months after infection (19–21). It can also not be excluded that re-infection with the same coronavirus type can occur within a single year (22). How the antibody titers translate to T cell immunity is not known. For influenza it is clear that recurrent infections and epidemics are due to the accumulation of mutations in the hemagglutinin and neuraminidase (23). However, the genetic drift of endemic coronaviruses seems to be considerably slower than for Influenza A and B (24,25). Collectively, this indicates that other frequently encountered pathogens besides the endemic coronaviruses could have generated the preexisting immunity.

In the present report, we evaluate commonly occurring human pathogens for epitopes with a very high similarity to potential SARS-CoV-2 epitopes.

## Results

We identified a number of pathogens commonly causing infections in the European population (Supplementary Tables 2-5). The list of pathogens included 32 viruses, 11 fungi, 26 bacteria and 2 parasites. We obtained all protein sequences for these pathogens from NCBI, and compared these to predicted HLA-I and HLA-II binding epitopes in SARS-CoV-2, and ranked the pathogens based on a relevance score (see Methods; Figure 1a).

**Figure 1.**
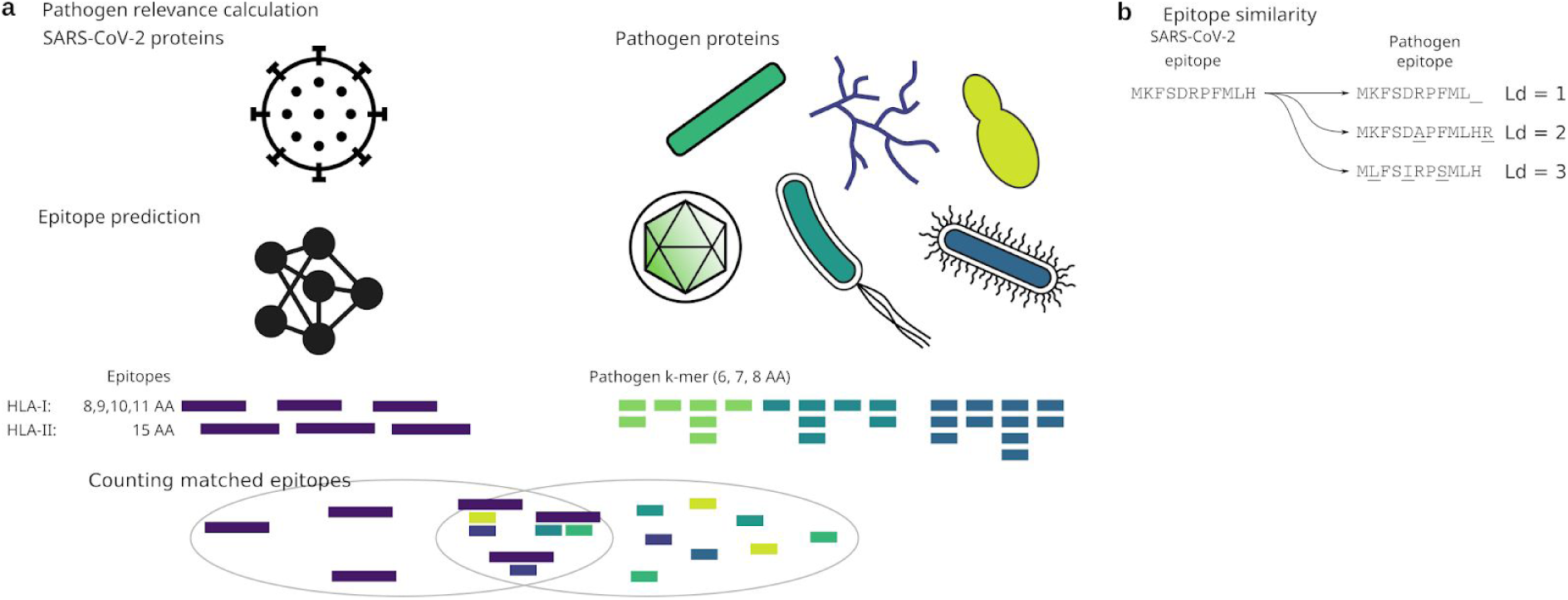
Analysis approach. a) 6, 7, 8mer k-mers were extracted from the proteins of relevant human pathogens and compared to epitopes predicted in the SARS-CoV-2 proteins. The epitope prediction was by netMHCpan and netMHCIIpan. Pathogens were ranked based on k-mer hits to the epitopes. b) Principle of Levenshtein distance of epitopes.

We observed different relevance scores for the same pathogen, depending on the length of *k*-mers (Supplementary Figure 1 and 2). For *k*=6 we found that all viruses were in the upper half when ranking the relevance scores, while the viruses were in the lower half at *k*=8. This was independent of the HLA-class epitopes (Supplementary Figure 1 and 2).

Given the overall similarity between coronaviruses, the endemic coronaviruses are expected to have the highest relevance score. Given the compact genome size and highly optimized proteins (26) it seems likely that only short stretches will have an exact match, even when using a reduced alphabet. Conversely, more complex organisms have larger and less optimized genes, why the probability of matching longer stretches increases. We therefore focus on pathogens with short (*k*=6) matches.

Within the top 10 ranking pathogens for HLA-I binding SARS-CoV-2 epitopes based on *k*=6, we find the two fungi Candida tropicalis and Cryptococcus neoformans, and the parasite Trichomonas vaginalis. Apart from the endemic coronaviruses HKU1, OC43, 229E, NL63 we find the double-stranded DNA virus Human alphaherpesvirus 3 (varicella-zoster virus, VZV), the double-stranded RNA virus Rotavirus A (RV), and the single-stranded negative RNA virus Influenza B (Figure 2a). When scoring the relevance based on matches to predi HLA-II SARS-CoV-2 epitopes, we find a similar picture, with the exception of the appearance of Human Gammaherpesvirus 4 (Epstein-Barr virus, EBV) in place of Trichomonas vaginalis (Figure 2b).

**Figure 2.**
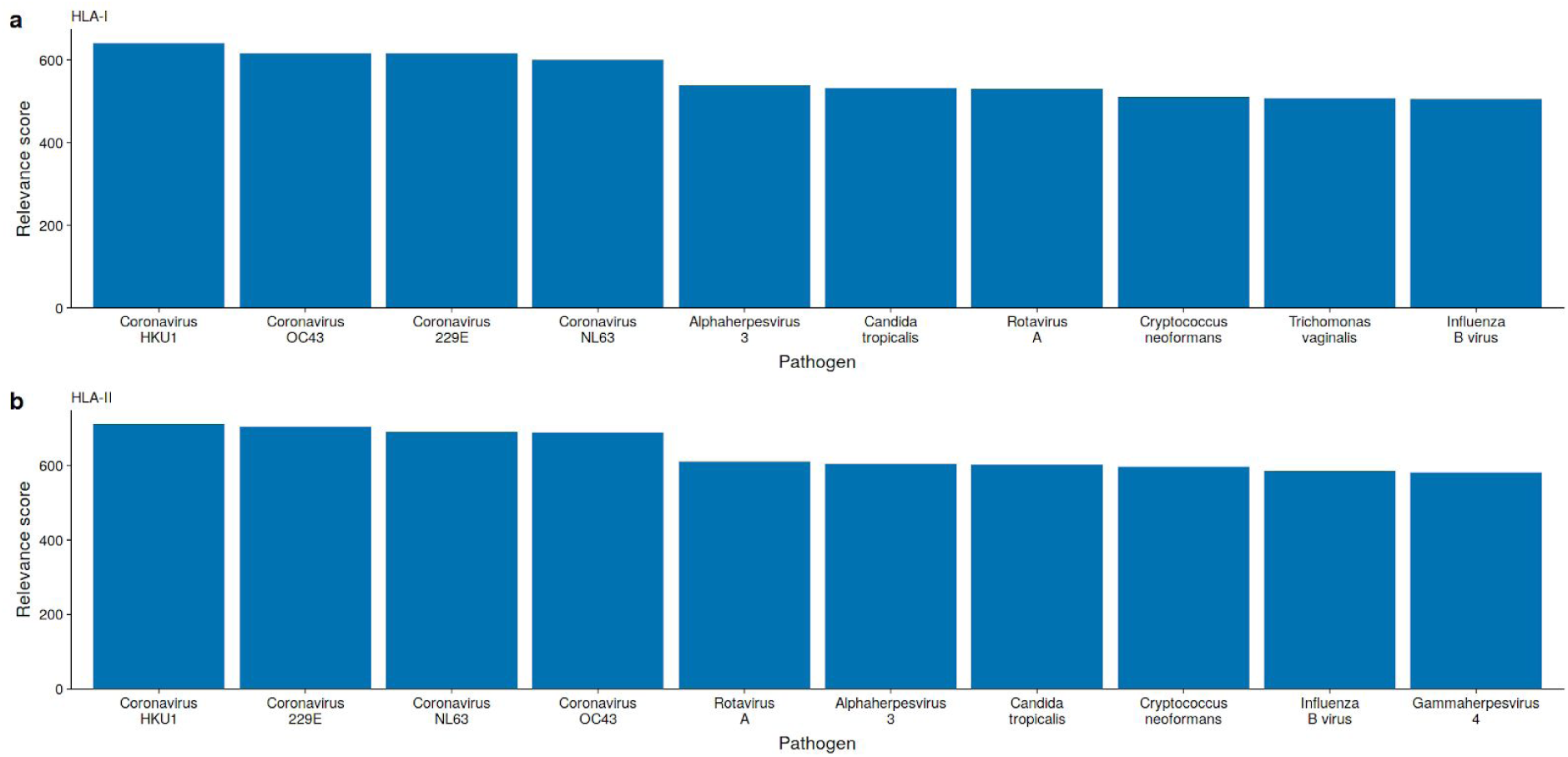
Some viruses and bacteria have peptides (k-mers) matching SARS-CoV-2 epitopes. The top 10 pathogen relevance score was averaged over *k*-mers at the length of 6, 7, and 8 amino acids for a) HLA-I, and b) HLA-II. Pathogen relevance score for each pathogen an *k* are presented in Supplementary figure 1 and 2.

Viral genes can be expressed at different times during an infection (27). However, during the multiplication phase of the virus, the viral products are expressed in excess. To avoid selection of pathogens based on highly similar, but rarely or poorly expressed proteins, we therefore focus on the viruses in the following analysis.

Using netMHCpan and netMHCIIpan we predicted the epitopes in the reference sequences for the coronaviruses OC43, HKU1, 229E, and NL63, as well as Influenza B, EBV, RV, and VZV. Assessing the similarity to SARS-CoV-2 predicted epitopes, we calculated the Levenshtein distance from each SARS-CoV-2 epitope to each of the predicted epitopes in the selected pathogens (Figure 1b). The Levenshtein edit distance accounts for addition, deletion, and exchange of amino acids to transform one epitope into another. We opted for this distance metric to allow differences in epitope lengths. The edit distance was calculated per analyzed HLA.

We found that the beta-coronaviruses OC43 and HKU1 had the highest number of epitopes identical to the predicted SARS-CoV-2 epitopes. For HLA-I bound epitopes we found 211 and 195 identical epitopes in HKU1 and OC3, respectively (Figure 3a). For HLA-II bound epitopes we found 493 and 464 identical epitopes in OC43 and HKU1, respectively (Figure 3b). When the similarity threshold was relaxed to an edit distance of 1 or 2 we found a similar pattern (Figure 3). Interestingly, allowing three insertions, deletions, or exchanges we find the highest number of similar SARS-CoV-2 HLA-I epitopes in VZV, followed by OC43, and HKU1 with 1292, 1189, and 1163 epitopes, respectively (Figure 3a). This was not reflected in HLA-II bound epitopes. This strong occurrence of SARS-CoV-2 similar VZV epitopes was mainly driven by epitopes on HLA-B and HLA-C, and to a minor degree on HLA-A (Supplementary figures 3-5).

**Figure 3.**
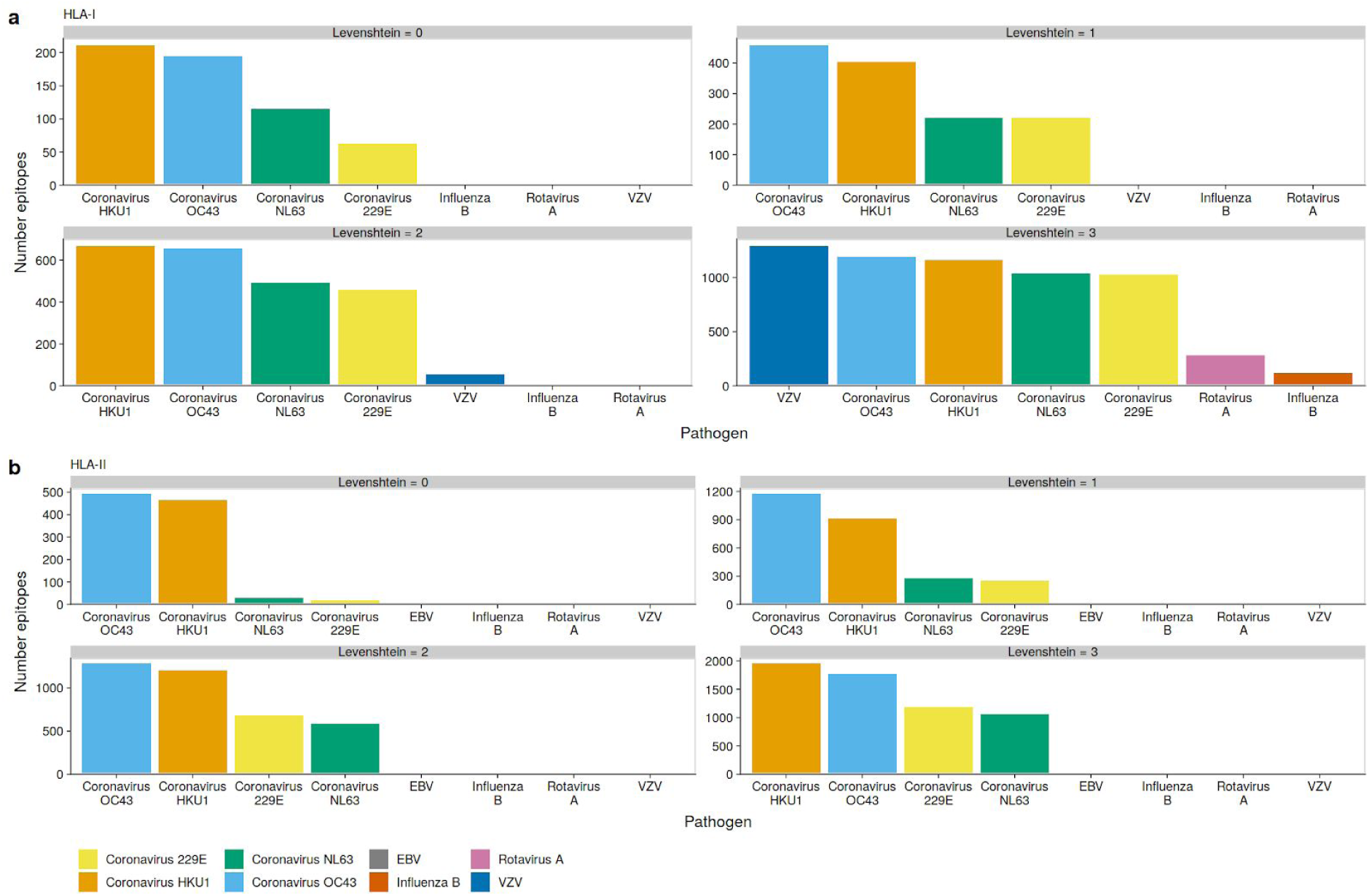
OC43 and HKU1 epitopes can be presented on many HLAs. Epitopes were predicted using netMHCpan and netMHCIIpan in selected pathogens. The similarity between each SARS-CoV-2 epitope and pathogen epitope was calculated using the Levenshtein distance, and the number of shortest matches was enumerated. a) Total number of HLA-I epitopes with an edit distance between 0 and 3. b) Total number of HLA-II epitopes with an edit distance between 0 and 3. The pathogens are ordered per plot from highest to lowest, while the fill colour is preserved.

## Discussion

There is mounting evidence for preformed immunity against the novel coronavirus SARS-CoV-2 (13–17). Reports on memory T cell response to the endemic coronaviruses are lacking, but since antibody titers appear short-lived and frequent re-infections cannot be excluded (19–22), it is probable that the endemic coronaviruses with low virulence do not create lasting immunity. This then begs the question, which pathogen can generate a memory T cell response cross reactive to SARS-CoV-2. Through *in silico* analysis of predicted SARS-CoV-2 epitopes and common pathogens we address this question.

When allowing some dissimilarity, VZV has the highest number of like SARS-CoV-2 epitopes. However, this we only find for HLA-I, and in particular HLA-B and HLA-C, but not HLA-II. Since SARS-CoV-2 specific T cells are found among both the HLA-I reactive CD8^+^ and among the HLA-II reactive CD4^+^ it is not likely that VZV is the pathogen behind the SARS-CoV-2 cross reactive T cells. Rather, only the endemic coronaviruses, and in particular the beta-coronaviruses, have the highest number of epitopes similar to those predicted in the SARS-CoV-2 virus.

While the prediction of HLA-I epitopes has high accuracy, the accuracy of HLA-II epitope prediction is lower than that for HLA-I (28). One reason for this lower accuracy of HLA-II epitope prediction is the dimer structure of the HLA-II making the peptide binding groove. In fact, the HLA-II α*β*-chain combinations have been estimated to be over 4000 in number. In contrast, the combination of the HLA-I and *β*2-microglobulin chains yields less combinatorial variation on the receptor. The epitope prediction with netMHCpan depends on a proper training set. By focusing on the most frequent european HLAs, inaccuracies in the epitope prediction are reduced or even avoided. We did not utilize epitope databases like VDJDB (29) and IEDB (30), the reason being that these databases have a natural bias towards laboratory model antigen. For instance, the most frequent epitopes in the Immune Epitope Database are human auto antigen and epitopes derived from Trypanosoma cruzi.

The amino acid set was reduced from the 20 standard amino acids to 15 amino acids by combining molecules with similar hydrophobic side chains. The advantage is the ease of sequence comparison without alignment since the *k*-mer is a complete substring of the target protein. While the reduced amino acid alphabet is derived from structural considerations (31), it cannot be excluded that alphabet reduction might skew the results. However, given the requirements of anchor amino acids, which are buried within the HLA-molecule, the final results should not change.

One limitation to the study is that we do not consider the frequency of pathogens nor the potential expression level of the proteins and accessibility for the immune system. Both arguably have an effect on the likelihood that an epitope can raise a robust immune response. However, both variables can only be assessed with great uncertainty. Another limitation to the study is the reliance on sequence similarity; although related epitopes are likely to interact with the same T cell receptor, this is not guaranteed (32).

The finding that the beta-coronaviruses OC43 and HKU1 are the most likely pathogens to give rise to SARS-CoV-2 preformed immunity means that the endemic coronaviruses must be able to generally raise a long lasting T cells response. Otherwise the epitopes recognized by the cross reactive T cells share no high degree of similarity.

In conclusion, we conjecture that other beta-coronaviruses are the source for SARS-CoV-2 cross reactive T cells.

## Methods

The SARS-CoV2 protein sequences were downloaded from ViralZone (33; https://viralzone.expasy.org/89966), accessed May 29, 2020. Uniprot IDs and common names are listed in Supplementary Table 1. The common human pathogens evaluated in this study are listed in Supplementary Tables 2-5. Pathogen protein sequences were extracted from the NCBI “non-redundant” protein database (“nr”, version 5, downloaded from ftp://ftp.ncbi.nlm.nih.gov/blast/db/FASTA on May 31, 2020). The extraction was per pathogen name, as stored in the Taxonomy database. For epitope comparison, the protein reference sequences for the coronaviruses OC43, HKU1, 229E, and NL63, and Influenza B, Human Gammaherpesvirus 4, Rotavirus A, and Human alphaherpesvirus 3, were downloaded from https://ftp.ncbi.nlm.nih.gov/refseq/release/viral on 26.Jun.2020.

All potential MHC class I and class II epitopes in the SARS-CoV2 protein sequences, and selected pathogen reference sequences, were identified using netMHCpan version 4.1 and netMHCIIpan version 4.0, respectively, with default settings (34). For this step, the evaluated HLAs listed in Supplementary Tables 6 and 7 were selected based on frequency in the European population as reported in the Allele Frequency Net Database (35; http://www.allelefrequencies.net, June 1, 2020).

The Levenshtein distance between each predicted SARS-CoV-2 epitope to a predicted epitope in a selected pathogen was calculated. The calculation as per pathogen and HLA and only the best match was used.

Short peptides of length *k (k*-mers) of lengths 6, 7, and 8 were extracted from the proteins of each of the relevant pathogens and counted, for each value of *k* separately Each such multiset of *k*-mers was prefiltered so that *k*-mers found only once in a pathogen were removed if those *k*-mers constituted a small fraction *f* or less of all *k*-mers found for the pathogen. The analysis was done separately for *f* in {0%, 1%, 5%}. The resulting *k*-mers were matched to the *k*-mers from the predicted epitopes for SARS-CoV-2 using a reduced 15 letter amino acid alphabet (31). This amino acid alphabet allows mismatches of amino acids with similar properties by letting the large hydrophobic amino acids V, L, I, and M be represented by L, the amino acids Y and F, which are hydrophobic with aromatic side chains by F, and the positively charged K and R by K.

The more complex an organism, the more proteins are expressed. This means that the probability of finding *k*-mers matching epitopes from SARS-CoV-2 increases. To overcome this problem we developed a *pathogen relevance score* to be evaluated for each triple of SARS-CoV-2 protein *p*, pathogen species *s* and value of *k* (note that the quantity also depends on the filter threshold f described above and on the considered set of epitopes, i.e. HLA class I or class II only or combined). We thus define

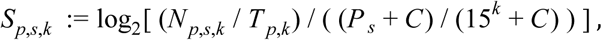

where *N*_*p,s,k*_ is the number of *k*-mer matched epitopes of protein *p* in species *s*, and 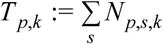 is the number of such epitopes of protein *p* across all species; their ratio is viewed in relation to the “richness” of the *k*-mer set of the pathogen species *s*, i.e. *P*_*s*_, the number of distinct *k*-mers in pathogen s, divided by the total number of possible k-mers over the reduced alphabet (15^*k*^). To avoid bias in favor of underrepresented species (very small *P*_*s*_), we add a regularizing constant C when computing the fraction of *k*-mers used by the species. Thus the score is a log-observed-vs-expected-ratio indicating whether species *s* matches unproportionally many SARS-CoV-2 epitopes that cannot be explained by its proteome size alone.

To focus on possibly relevant pathogens in general rather than on a single SARS-CoV-2 protein, we also considered an aggregated score

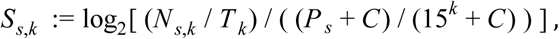

where *N*_*s,k*_ is the total number of *k*-mer matched epitopes in species *s*, and 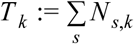 is the number of such epitopes across all species. The pathogens were then ranked according to score (the highest score obtaining rank 1; the lowest score rank *n*, where *n* is the number of considered species). Then the average rank is computed over the different parameter combinations k=6,7,8 and f=1%.

## Acknowledgements

This work was supported by grants from the Mercator Foundation (St-2018-0014), BMBF e:KID (01ZX1612A), and BMBF NoChro (FKZ 13GW0338B).

## Author contributions

Conceptualization: U.S. and T.R.; Data curation: U.S. and S.R.; Formal analysis: U.S. and S.R.; Funding acquisition: S.R. and N.B.; Investigation: U.S. and S.R.; Methodology: S.R.; Resources: T.H.W.; Visualization: U.S.; Writing – original draft: U.S. and S.R.

