## Supplement Information for "SARS-CoV-2 reactive T cells in uninfected individuals are likely expanded by beta-coronaviruses"

* Equal contribution

### Supplementary Methods

#### Workflow and Software

Our workflow is available as a Snakefile for use with Snakemake (1), starting with the download of the "nr" database from NCBI, at gitlab at the URL https://gitlab.com/svenrahmann/corona. The following steps are included, each performed by a specific Python script contained in the above repository.

Required inputs to the workflow are:

- a FASTA file with SARS-CoV-2 protein sequences (sara-cov2.fa),
- a tab-separated file with selected taxIDs of species to consider (species_taxid.tsv),
- a text file with predicted epitopes of SARS-CoV-2 (sars_mhci_peptides.txt, sars_mhcii_peptides.txt; one peptide per line)

All of these are provided in the repository.

The workflow consists of the following steps.

1. Download of the NCBI nr database: Snakefile (rule get_nr)
2. Extract protein sequences of common human pathogens from nr: Snakefile (rule select_species); filterspecies.py
3. Compute statistics on the number of sequences of each selected pathogen present in the nr database: Snakefile (rule count_selected)
4. Extract k-mers from each species' protein sequences for several values of k and for several reduced alphabets: Snakefile (rule get_kmers); getcounters.py
5. Filter out k-mers that only occur once in the set of all proteins of a species for different k, alphabets and filter thresholds: Snakefile (rule filter_kmers): filterkmers.py
6. Create a table of SARS-CoV-2 epitopes by protein with k-mer matches against pathogens. The resulting output files (results/epitopes_{class}_{k}_{alphabet}_{filterthreshold}) have a tab-separated tabular format with 3 columns: (SARS-CoV-2 protein name, pathogen name, peptide sequence of matched SARS-CoV-2 epitope). After all epitopes matched by the same pathogen have been shown, a statistics line (starting with '#') is inserted, giving the pathogen relevance score and other statistical information. At the end of the file, several summary statistics (over all proteins, per pathogen) are given (lines starting with '+'), including overall pathogen relevance scores. The same information is also written into a summary file: Snakefile (rules show_epitopes; summarize_epitopes); showepitopes.py.
7. Compute average pathogen ranks from different values of k and filter thresholds, for each epitope class I and II separately and combined, and generate plots of ranked ranks: Snakefile (rule aggregate_results); aggregate.py

### Supplementary Tables

#### Supplementary Table 1: SARS-CoV-2 proteins

Name and function of the SARS-CoV2 proteins obtained from ViralZone (2)

| UniProt ID | Common name |
|---|---|
| P0DTC1 | Replicase polyprotein 1a (pp1a) |
| P0DTD1 | Replicase polyprotein 1ab (pp1ab) |
| P0DTC2 | Spike glycoprotein (S) |
| P0DTC3 | ORF3a protein (NS3a) |
| P0DTC4 | Envelope small membrane protein (E) |
| P0DTC5 | Membrane protein (M) |
| P0DTC6 | ORF6 protein |
| P0DTC7 | ORF7a protein |
| P0DTD8 | ORF7b protein |
| P0DTC8 | ORF8 protein |
| P0DTC9 | Nucleoprotein (N) |
| P0DTD2 | ORF9b protein |
| P0DTD3 | ORF14 protein |

### Supplementary Table 2: Selected viruses

Names and taxIDs obtained from the NCBI Taxonomy database (3).

| Family | Scientific name | TaxID | Taxon level |
|---|---|---|---|
| **ssRNA(+)** | | | |
| Caliciviridae | Norwalk virus | 11983 | Species |
| Coronaviridae | Human coronavirus OC43 | 31631 | Species |
| | Human coronavirus HKU1 | 290028 | Species |
| | Human coronavirus 229E | 11137 | Species |
| | Human coronavirus NL63 | 277944 | Species |
| Matonaviridae | Rubella virus | 11041 | Species |
| Picornaviridae | Human rhinovirus A* | 147711 | Species |
| | Human rhinovirus B* | 147712 | Species |
| | Human rhinovirus C* | 463676 | Species |
| | Enterovirus B | 138949 | Species |
| | Human hepatitis A virus | 208726 | Subtype |
| | Hepatitis B virus | 10407 | Species |
| | Hepatitis E virus | 12461 | no rank |
| | Hepacivirus C | 11103 | Species |
| | Human poliovirus 1 | 12080 | Subtype |
| | Human poliovirus 2 | 12083 | Subtype |
| | Human poliovirus 3 | 12086 | Subtype |
| **ssRNA(-)** | | | |
| Orthomyxoviridae | Influenza A virus | 11320 | Species |
| | Influenza B virus | 11520 | Species |
| Paramyxoviridae | Human respirovirus 1 | 12730 | Species |
| | Human respirovirus 3 | 11216 | Species |
| | Human rubulavirus 2 | 1979160 | Species |
| | Human rubulavirus 4 | 1979161 | Species |
| | Mumps rubulavirus | 1979165 | Species |
| | Measles morbillivirus | 11234 | Species |
| Pneumoviridae | Human metapneumovirus | 162145 | Species |
| **dsRNA** | | | |
| Reoviridae | Rotavirus A | 28875 | Species |
| **dsDNA** | | | |
| Herpesviridae[1] | Human alphaherpesvirus 1 | 10298 | Species |
| | Human alphaherpesvirus 3 | 10335 | Species |

| | | | |
|---|---|---|---|
| | Human gammaherpesvirus 4 | 10376 | Species |
| | Human betaherpesvirus 5 | 10359 | Species |
| Papillomaviridae | Human papillomavirus | 10566 | Species |

[*]Subspecies were included by necessity.
[1]Common names: Human alphaherpesvirus 1: Herpes simplex virus type 1; Human alphaherpesvirus 3: Varicella-zoster virus; Human gammaherpesvirus 4: Epstein-Barr virus; Human betaherpesvirus 5: Human cytomegalovirus

### Supplementary Table 3. Selected fungi

Names and taxIDs obtained from the NCBI Taxonomy database (3)

| Genus | Scientific name | TaxID | Taxon level |
|---|---|---|---|
| **Yeast** | | | |
| Candida | Candida albicans | 5476 | Species |
| | Candida glabrata | 5478 | Species |
| | Candida tropicalis | 5482 | Species |
| **Fungus** | | | |
| Aspergillus | Aspergillus fumigatus | 746128 | Species |
| | Aspergillus flavus | 5059 | Species |
| | Aspergillus niger | 5061 | Species |
| Cryptococcus | Cryptococcus neoformans | 5207 | Species |
| Pneumocystis | Pneumocystis jirovecii | 42068 | Species |
| Stachybotrys | Stachybotrys chartarum* | 74722 | Species |
| Trichophyton | Trichophyton rubrum | 5551 | Species |
| | Trichophyton mentagrophytes | 523103 | Species |

*Subspecies were included by necessity

### Supplementary Table 4. Selected bacteria

Names and taxIDs obtained from the NCBI Taxonomy database (3)

| Genus | Scientific name | TaxID | Taxon level |
|---|---|---|---|
| **Atypical** | | | |
| Mycoplasma | Mycoplasma pneumoniae | 2104 | Species |
| | Mycoplasma genitalium | 2097 | Species |
| **Gram-negative** | | | |
| Bordetella | Bordetella pertussis | 520 | Species |
| Burkholderia | Burkholderia cepacia | 292 | Species |
| Campylobacter | Campylobacter jejuni | 197 | Species |
| Chlamydia | Chlamydia trachomatis | 813 | Species |
| Escherichia | Escherichia coli | 562 | Species |
| Haemophilus | Haemophilus influenzae | 727 | Species |
| Klebsiella | Klebsiella pneumoniae | 573 | Species |
| Legionella | Legionella pneumophila | 446 | Species |
| Neisseria | Neisseria gonorrhoeae | 485 | Species |
| Proteus | Proteus mirabilis | 584 | Species |
| Pseudomonas | Pseudomonas aeruginosa | 287 | Species |
| Salmonella | Salmonella enterica subsp. enterica serovar Typhimurium | 90371 | Serovar |
| | Salmonella enterica subsp. enterica serovar Enteritidis | 149539 | Serovar |
| Shigella | Shigella flexneri | 623 | Species |
| | Shigella boydii | 621 | Species |
| | Shigella dysenteriae* | 622 | Species |
| **Gram-positive** | | | |
| Bacillus | Bacillus cereus | 1396 | Species |
| Clostridium | Clostridium tetani | 1513 | Species |
| | Clostridium difficile* | 1496 | Species |
| Corynebacterium | Corynebacterium diphtheriae | 1717 | Species |
| Staphylococcus | Staphylococcus aureus | 1280 | Species |
| | Staphylococcus epidermidis | 1282 | Species |
| Streptococcus | Streptococcus pneumoniae | 1313 | Species |
| | Streptococcus pyogenes | 1314 | Species |

*Subspecies were included by necessity

### Supplementary Table 5. Selected parasites

Names and taxIDs obtained from the NCBI Taxonomy database (3)

| Genus | Scientific name | TaxID | Taxon level |
|---|---|---|---|
| **Parasites** | | | |
| Trichinella | Trichinella spiralis | 6334 | Species |
| Trichomonas | Trichomonas vaginalis | 5722 | Species |

### Supplement Table 6. Used HLA-I molecules

Allele frequencies for Europe were obtained from the Allele Frequency Net Database (4).

| Locus | Allele | % individuals with allele |
|---|---|---|
| **A** | | |
| | A*02:01 | 15.08 |
| | A*01:01 | 10.28 |
| | A*03:01 | 8.68 |
| | A*24:02 | 6.14 |
| | A*11:02 | 3.80 |
| **B** | | |
| | B*08:01 | 6.77 |
| | B*07:02 | 6.02 |
| | B*44:02 | 5.48 |
| | B*51:01 | 4.89 |
| | B*35:01 | 4.73 |
| **C** | | |
| | C*07:01 | 11.58 |
| | C*07:02 | 8.21 |
| | C*04:01 | 7.80 |
| | C*05:01 | 6.09 |
| | C*06:02 | 5.46 |

### Supplementary Table 7. Used HLA-II molecules

Allele frequencies for Europe were obtained from the Allele Frequency Net Database (4).

| Locus | Allele | % individuals with allele |
|---|---|---|
| **DPA1** | | |
| | DPA1*01:03 | 19.40 |
| | DPA1*02:01 | 6.60 |
| | DPA1*02:02 | 1.04 |
| | DPA1*04:01 | 0.60 |
| | DPA1*01:04 | 0.50 |
| **DPB1** | | |
| | DPB1*04:01 | 27.63 |
| | DPB1*02:01 | 13.17 |
| | DPB1*04:02 | 10.19 |
| | DPB1*03:01 | 7.60 |
| | DPB1*107:01 | 6.50 |
| **DQA1** | | |
| | DQA1*05:05 | 25.51 |
| | DQA1*05:01 | 12.92 |
| | DQA1*01:02 | 10.91 |
| | DQA1*03:01 | 8.25 |
| | DQA1*02:01 | 6.61 |
| **DQB1** | | |
| | DQB1*03:01 | 12.81 |
| | DQB1*02:01 | 9.67 |
| | DQB1*05:01 | 6.50 |
| | DQB1*06:02 | 6.17 |
| | DQB1*02:02 | 5.71 |

### Supplementary Figures

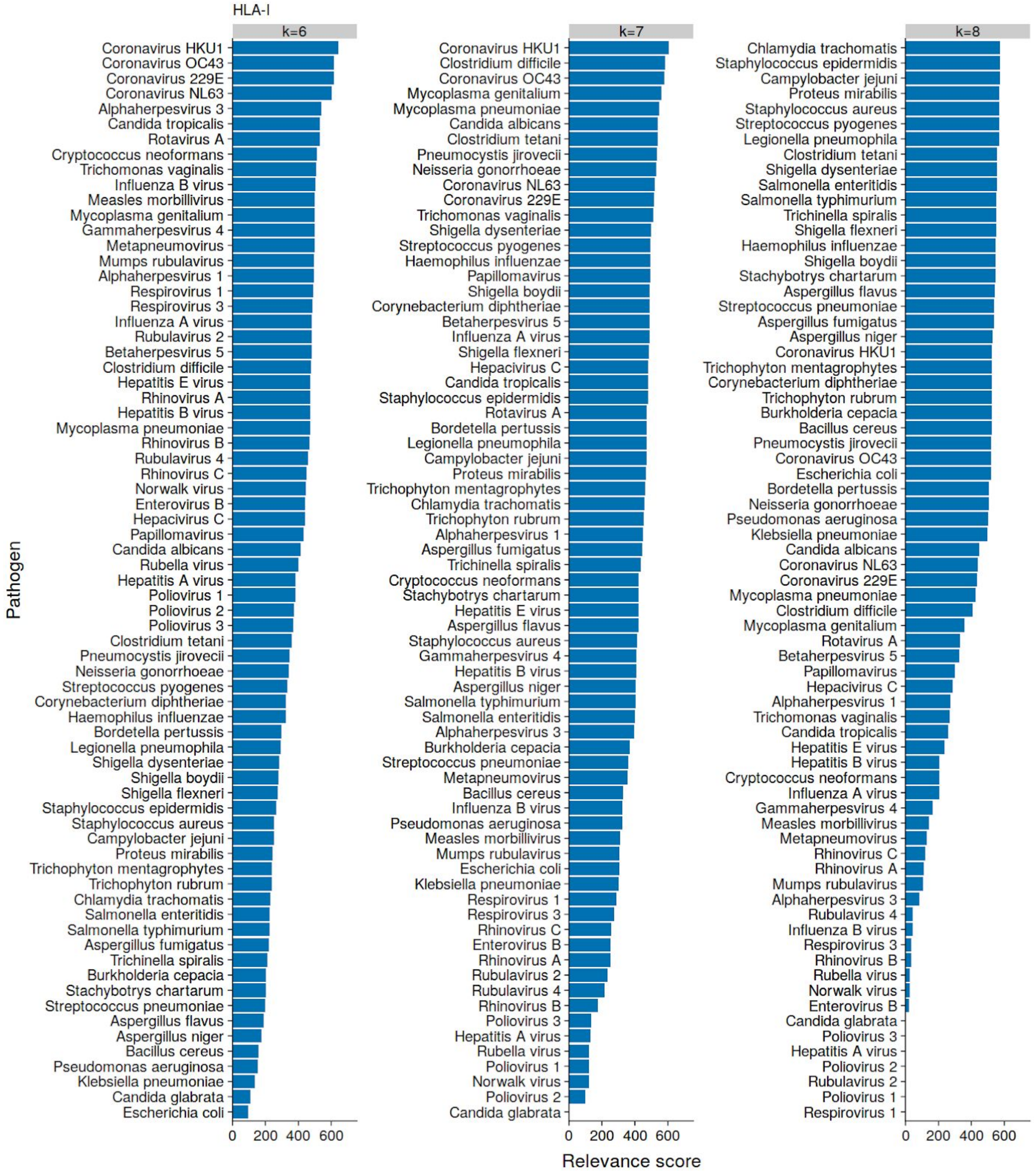


Supplementary Figure 1: Pathogen relevance score for HLA-I SARS-CoV-2 epitopes

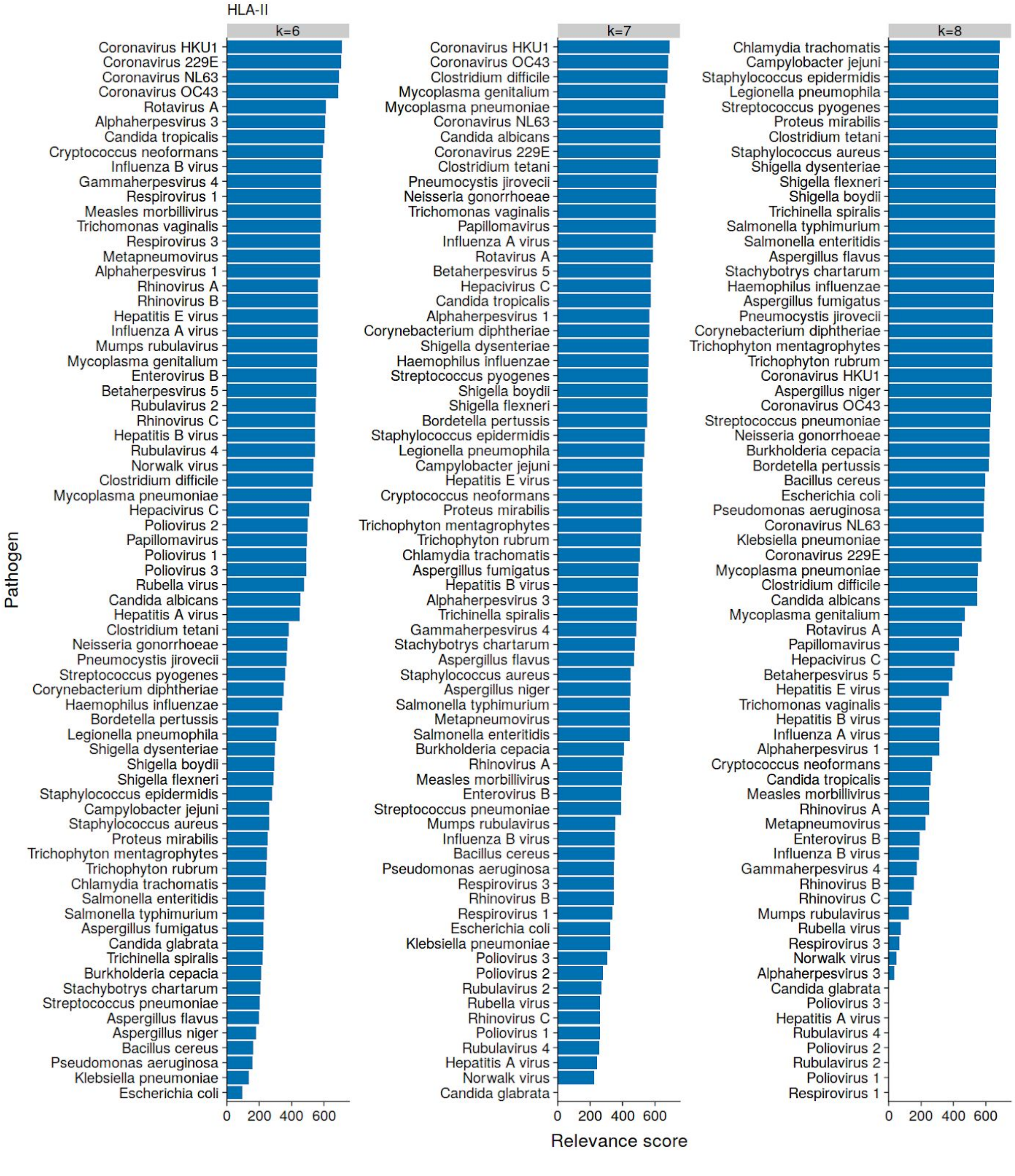


Supplementary Figure 2: Pathogen relevance score for HLA-II SARS-CoV-2 epitopes

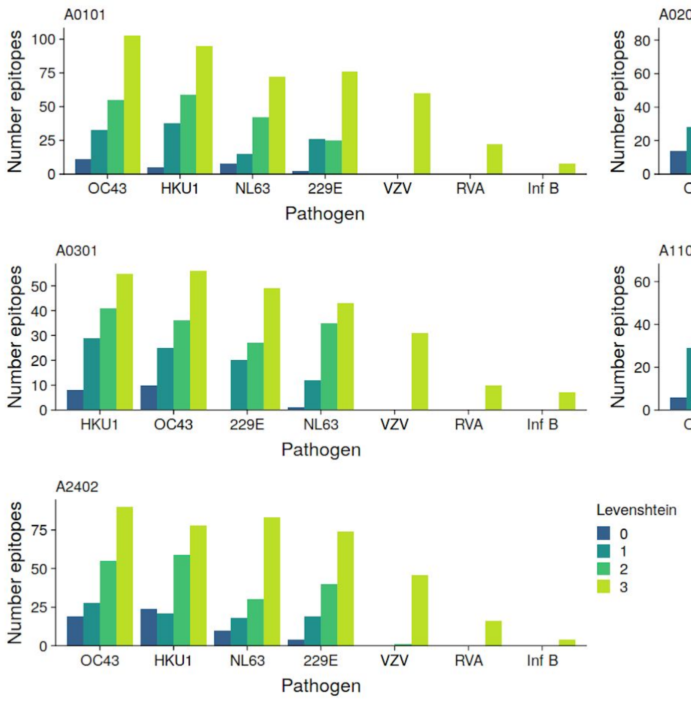


Supplementary Figure 3: Levenshtein distance between HLA-A SARS-CoV-2 epitopes and epitopes in selected pathogen

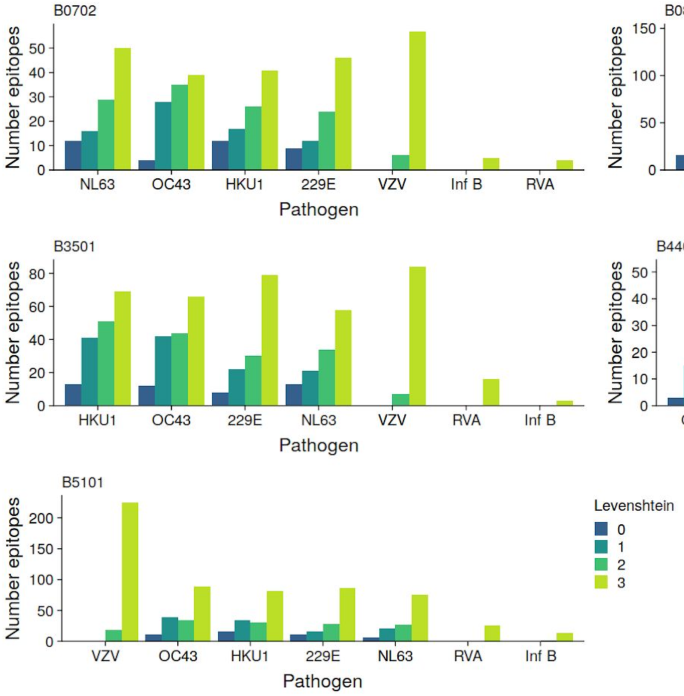


Supplementary Figure 4: Levenshtein distance between HLA-B SARS-CoV-2 epitopes and epitopes in selected pathogen

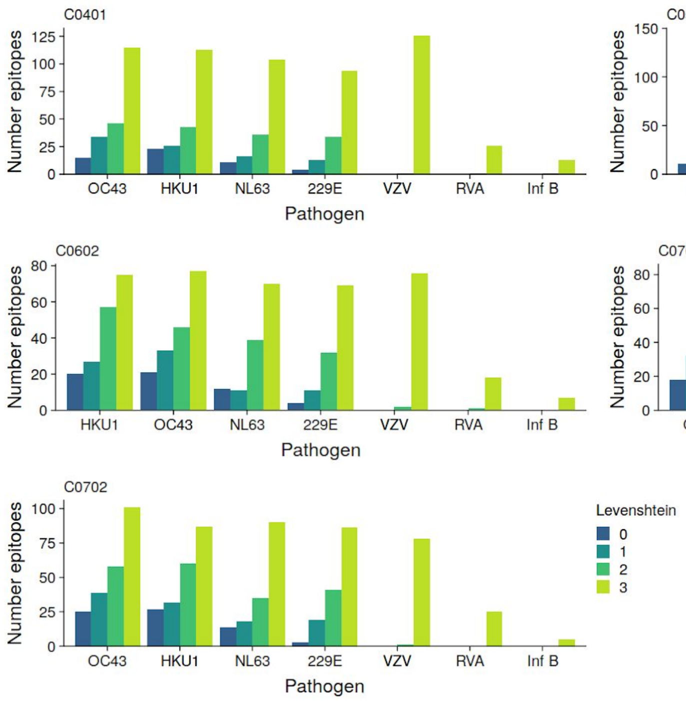


Supplementary Figure 5: Levenshtein distance between HLA-C SARS-CoV-2 epitopes and epitopes in selected pathogen
